## Supplemental Figures for "A second DNA binding site on RFC facilitates clamp loading at gapped or nicked DNA"

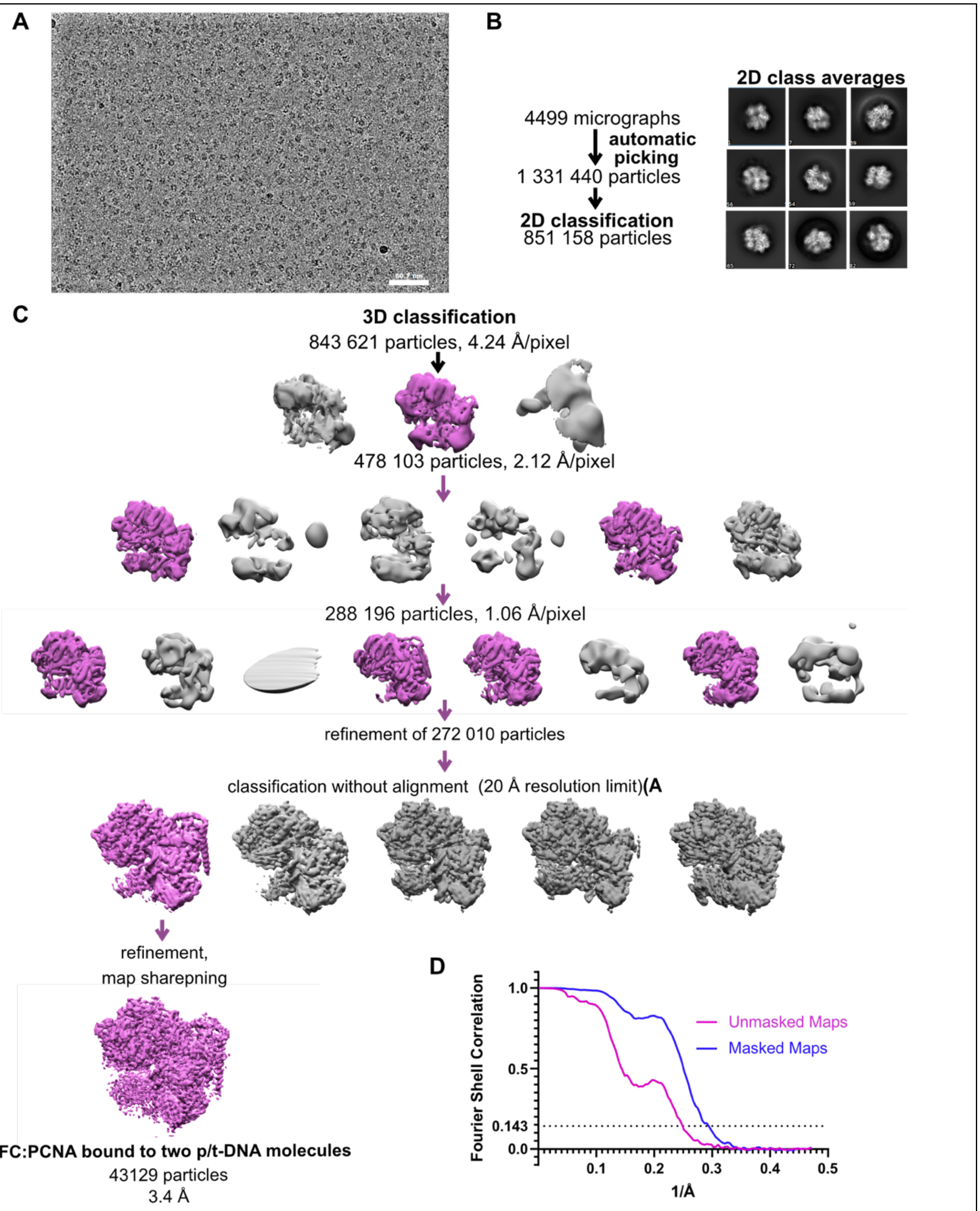


**Supplemental Figure 1.1. cryo-EM processing of RFC:PCNA in the presence of p/t DNA. (A)** Downfiltered micrograph taken on a Titan Krios with a Gatan K3 detector. **(B)** 2D class averages show well-resolved features and different views**.** **(C)** Data processing and 3D classification scheme, as described in Gaubitz et al. 2021. Here, a class showing RFC:PCNA bound to two p/t DNA molecules was further refined. The BRCT of Rfc1 domain is most visible in this class. **(D)** Fourier Shell correlation (FSC) curves obtained from postprocessing in Relion for the two halves of the unmasked and masked reconstructions.

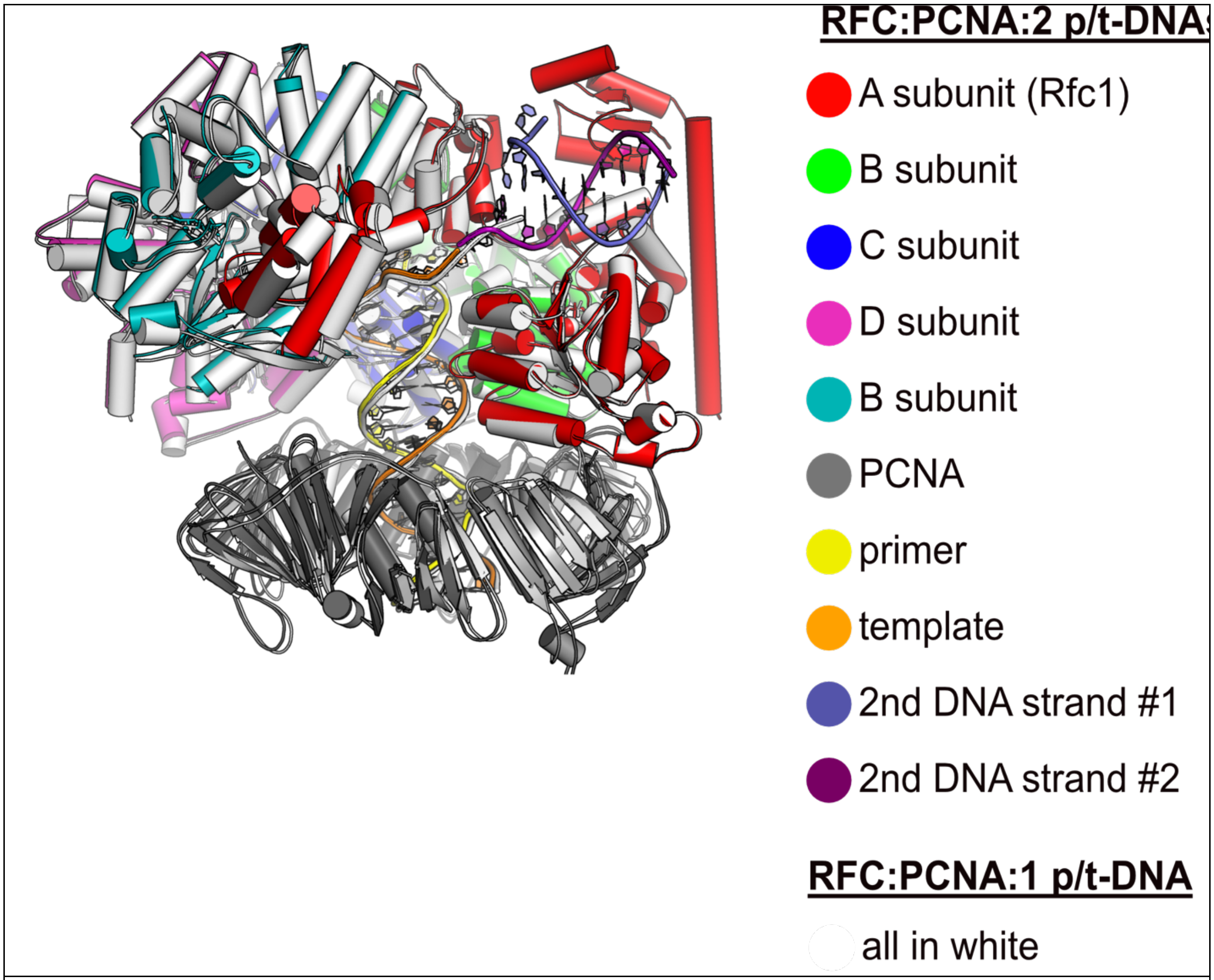


**Supplemental Figure 1.2. Structural similarity between structures with one or two p/t-DNAs bound.** The structure of RFC:PCNA bound to a single p/t-DNA (white; Gaubitz et al 2021) compared to the structure reported here bound to two p/t-DNAs (colored). The two structures were superposed only using the AAA+ module of RFC-A; the complex looks largely identical except for the presence of the BRCT domain, linker helix, and second DNA.

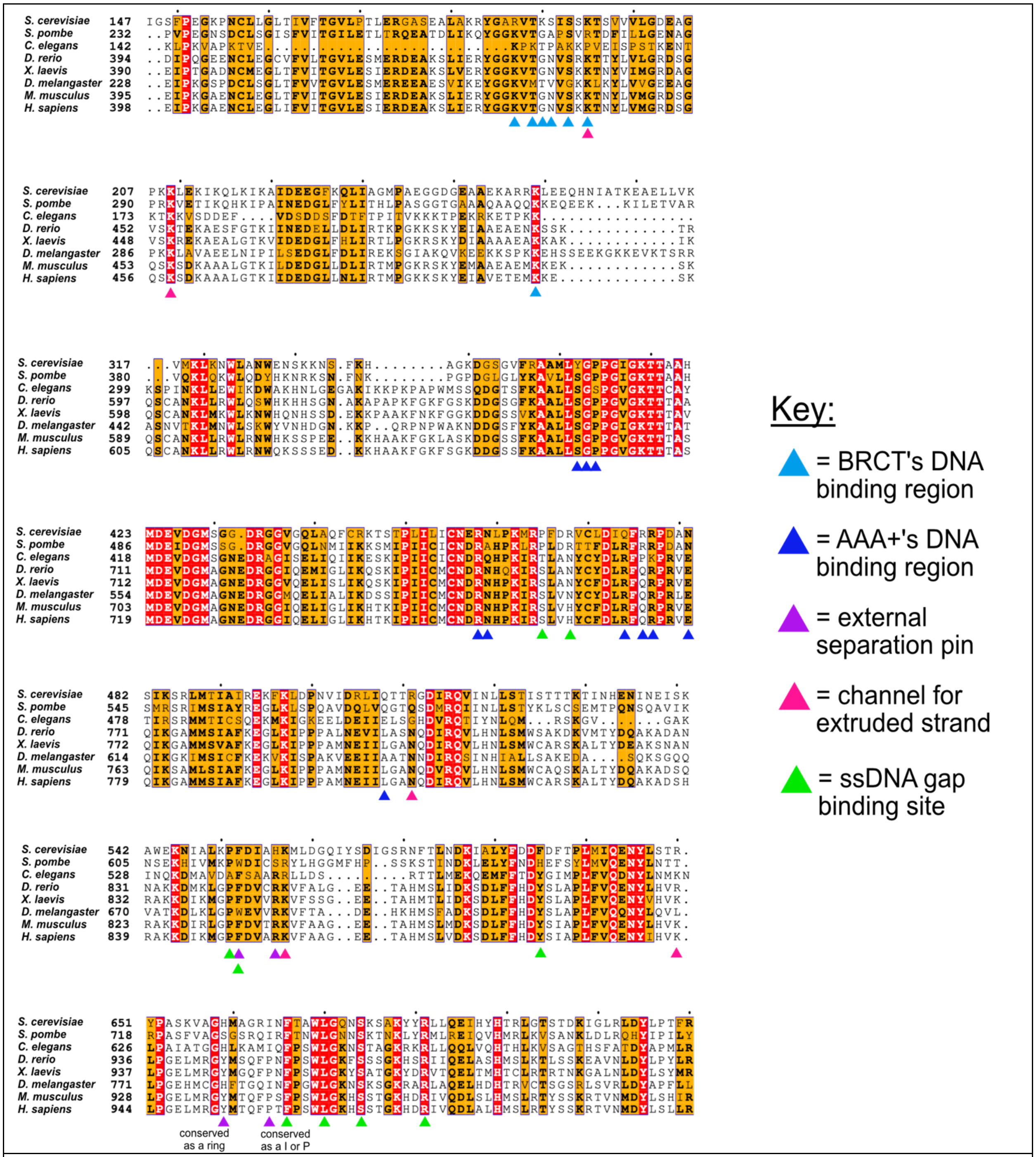


**Supplemental Figure 1.3: conservation of key residues for binding DNA.** The key residues in the two DNA binding sites of Rfc1 are conserved among eukaryotes. Sequence alignment shows the conservation of the 'separation pin' among 8 eukaryotic species. The conserved sequences are marked by blue boxes. The fully conserved residues are in white with a red background, the highly conserved residues are in black with a yellow background, and the less conserved ones are in black. The BRCT's and AAA+'s DNA binding residues are pointed out by light- and dark-blue arrows, respectively. The external separation pin residues are highlighted by purple arrows, and the residues in the channel for extruded strand are pointed out by pink arrows, and the residues for ssDNA gap binding site are pointed out by green arrows. Position 659 is conserved as a ring, and Position 664 as I or P.

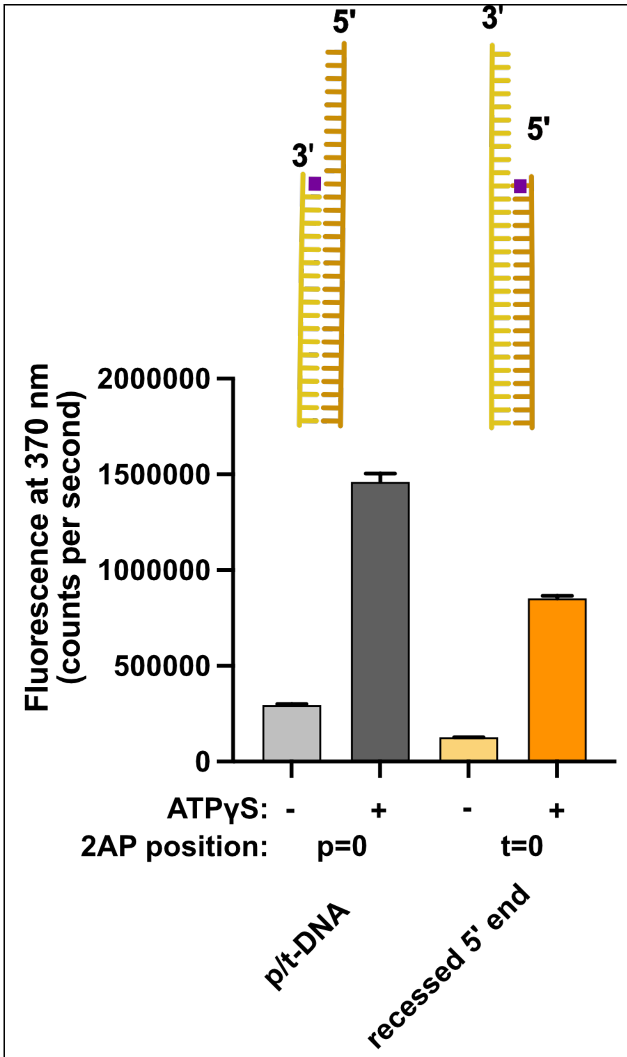


**Supplemental Figure 1.4: Binding of 5' recessed ends to RFC.** 2AP fluorescence was measured using DNA with a 3‘ recessed end or a 5' recessed end. The 2AP probe was placed at the 3‘ end of the primer or the 5' end of the template strand, respectively. In the presence of RFC and PCNA, we observe an ATP-analog dependent increase in 2AP fluorescence that is comparable to the increase seen with the well-characterized p/t-DNA substrate with 2AP at the 3' end of the primer strand. This result indicates that 5' recessed DNA can bind RFC and suggests that the 5' base is being melted. Because this increase is ATP-dependent and 5‘ recessed DNA does not stimulate ATP hydrolysis, this suggests that the melting does not occur at the internal separation pin, but occurs externally in the ATP-dependent BRCT-docked state.

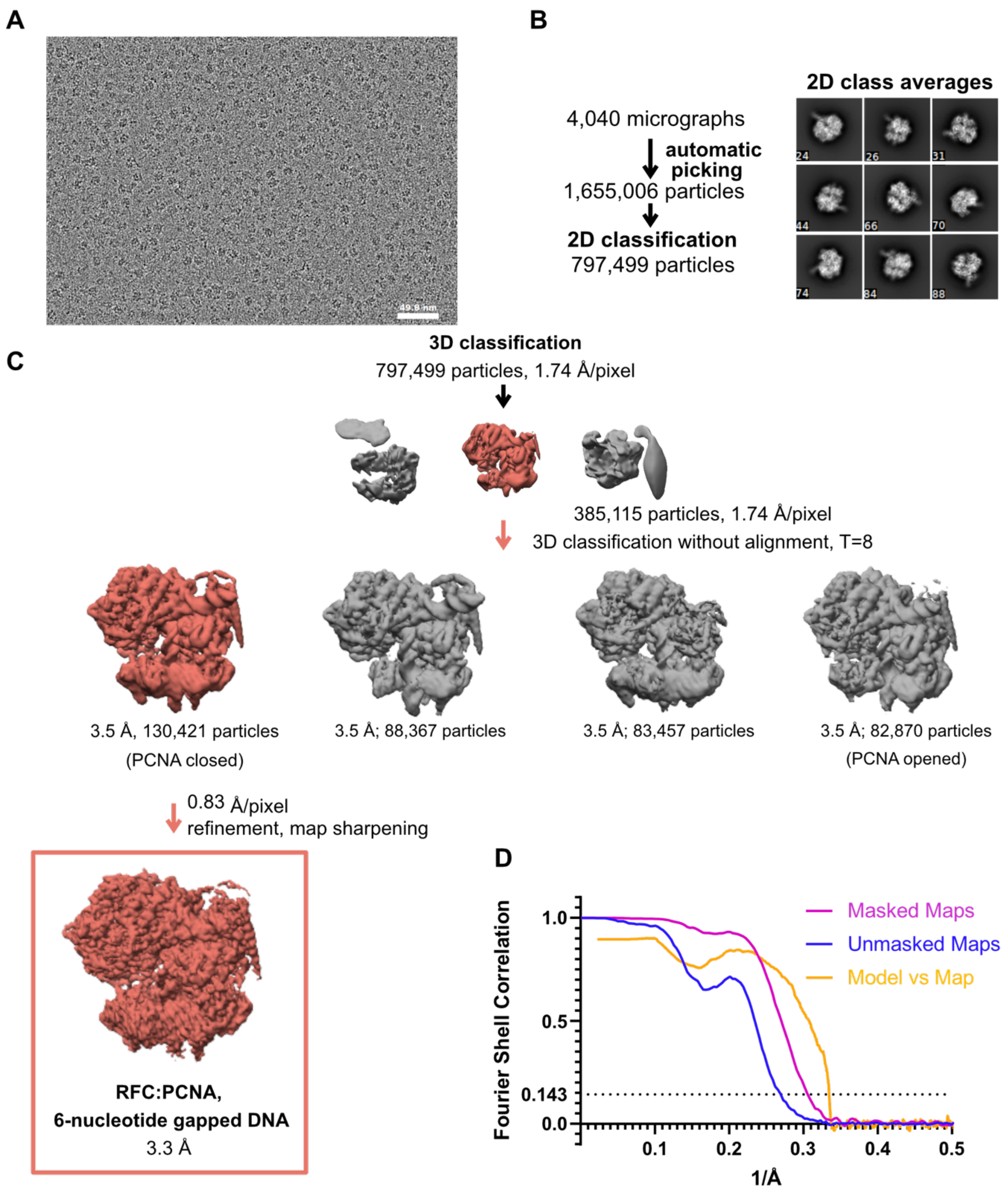


**Supplemental Figure 3.1: cryo-EM processing of RFC:PCNA in the pesence of dsDNA with a 6 nucleotide gap. (A)** Micrograph taken on a Talos Arctica with a Gatan K3 detector. **(B)** 2D class averages show well-resolved features and different side views**. (C)** The downfiltered reconstruction of RFC.PCNA bound to two p/t DNA molecules was used as a reference for 3D classification. **(D)** Fourier Shell correlation (FSC) curves obtained from postprocessing in Relion for the two halves of the unmasked and masked reconstructions as well as model vs. map curve.

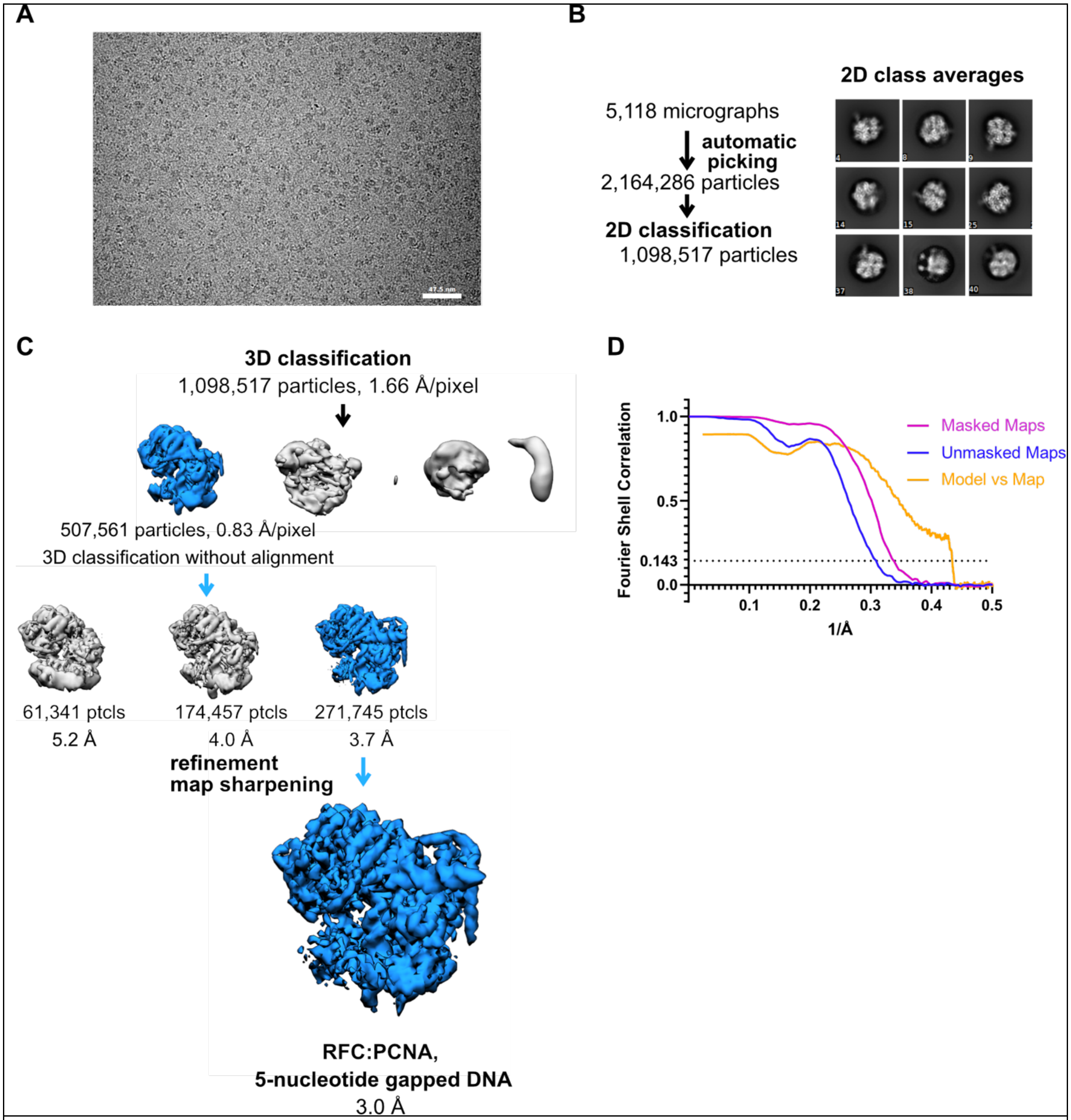


**Supplemental Figure 3.2. cryo-EM processing of RFC:PCNA in the presence of dsDNA with a 5 nucleotide gap. (A)** Micrograph taken on a Titan Krios with a Gatan K3 detector. **(B)** 2D class averages show different side views**. (C)** Data processing and 3D classification scheme. The cryo-EM reconstruction of RFC:PCNA bound to dsDNA with a 6 nucleotide gap was used as a 3D reference for 3D classification. **(D)** Fourier Shell correlation (FSC) curves obtained from postprocessing in Relion for the two halves of the unmasked and masked reconstructions, and model vs map curve.

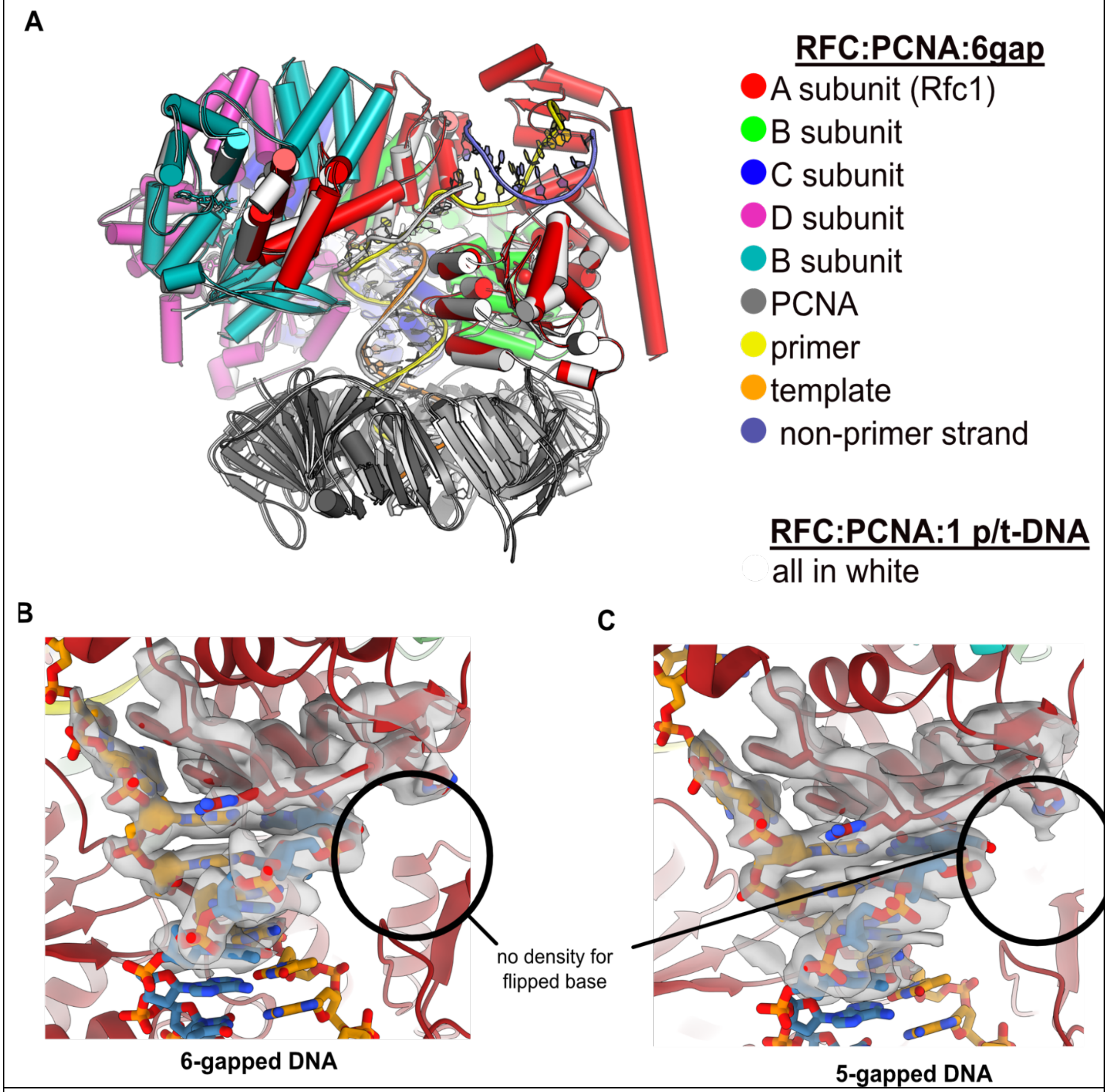


**Supplemental Figure 3.3. (A)** The ssDNA region in the structure of RFC:PCNA with dsDNA with a 6-nucleotide gap resembles the template overhang observed in RFC:PCNA bound to a single p/t-DNA.The structure of RFC:PCNA bound to a single p/t-DNA (white; Gaubitz et al 2021) compared to the structure reported here bound to 6 gapped DNA (colored). The two structures were superposed using the AAA+ module of RFC-A. The ssDNA region of the 6-gapped DNA exits RFC's central chamber through the A-gate, like the template overhang in the structure of RFC:PCNA bound to one p/t-DNA molecule. **(B&C)** RFC binds 5- or 6-gapped DNA without melting at the external separation pin. **(B)** Close-up view on the DNA at the external separation pin, cryo-EM density is shown in grey. The 6-gapped DNA binds to RFC:PCNA without melting at the external seperation pin (circled area). **(C)** The 5-gapped DNA also binds to RFC:PCNA without melting at the external seperation pin (circled area).

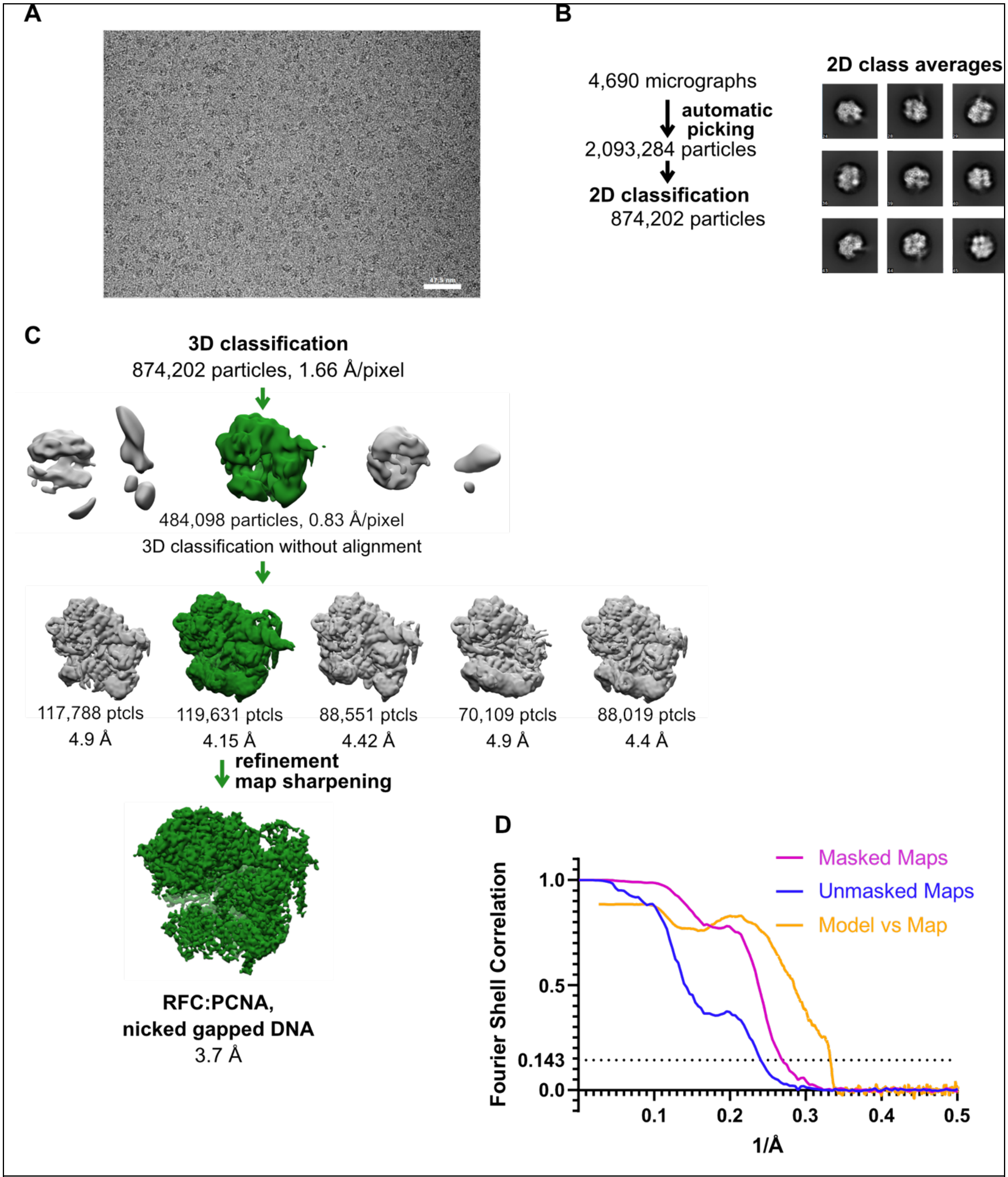


**Supplemental Figure 4.1: cryo-EM processing of RFC:PCNA in the presence of nicked dsDNA: (A)** Micrograph taken on a Titan Krios with a Gatan K3 detector. **(B)** 2D class averages show well-resolved features and different side views**. (C)** The reconstruction of RFC:PCNA bound to the 5-gapped dsDNA was downfiltered and used as a reference for 3D classification. **(D)** Fourier Shell correlation (FSC) curves obtained from postprocessing in Relion for the two halves of the unmasked and masked reconstructions as well as the model vs map curve.

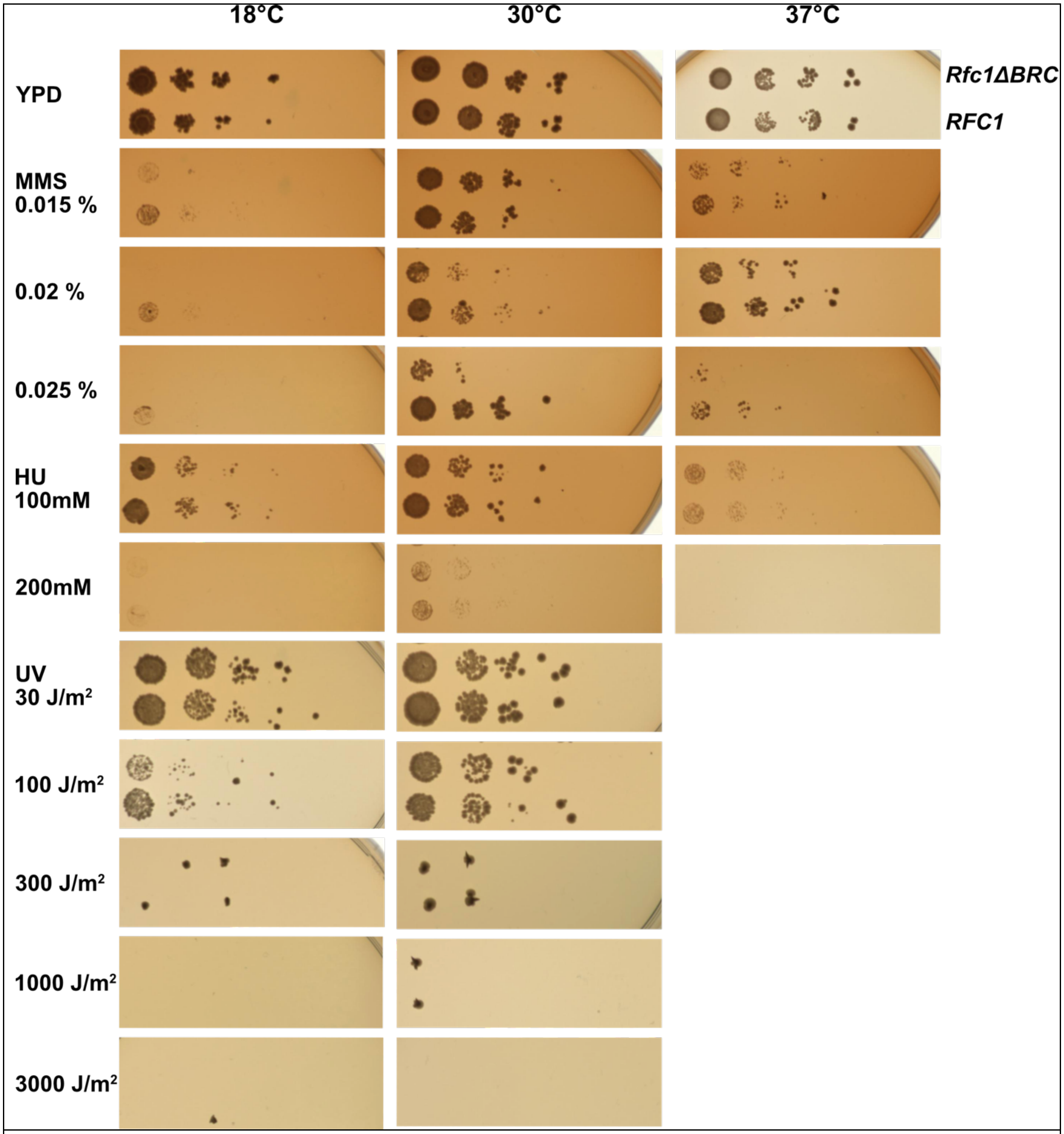


**Supplemental Figure 5.1. Deletion of the BRCT domain of Rfc1 results in a DNA damage repair defect.** Yeast carrying the sole copy of the RFC1 gene on a plasmid were subjected to various treatments that stress DNA metabolism. Multiple concentrations of the DNA damaging agents were used at different temperatures, including 0.015, 0.02, and 0.025% MMS, 100 and 200mM HU, at 18, 20, and 37°C, and 30, 100, 300, 1000, and 3000 J/m$^2$ UV at 18 and 30 °C. Rfc1-ΔBRCT yeast exhibit a growth defect only with the DNA alkylating agent methyl methanesulfonate (MMS), but not with hydroxyurea (HU) or ultraviolet radiation (UV). The phenotype is not dependent on temperature.

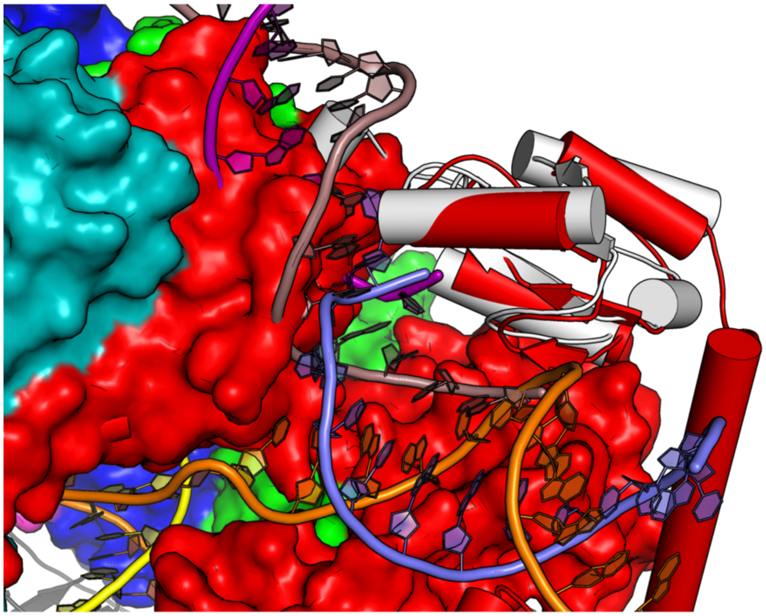


**Supplement Figure 6.1. DNA binding mode of free BRCT is incompatible with the conformation of open RFC.** The structure of free BRCT (PDB code: 2K7F, Kobayashi et al., 2010) bound to DNA compared to the structure of RFC:PCNA bound to nicked DNA. The structures were superposed using the BRCT. The DNA binding mode of free BRCT is incompatible with the conformation of RFC reported here. The DNA clashes with the collar domain of A subunit.

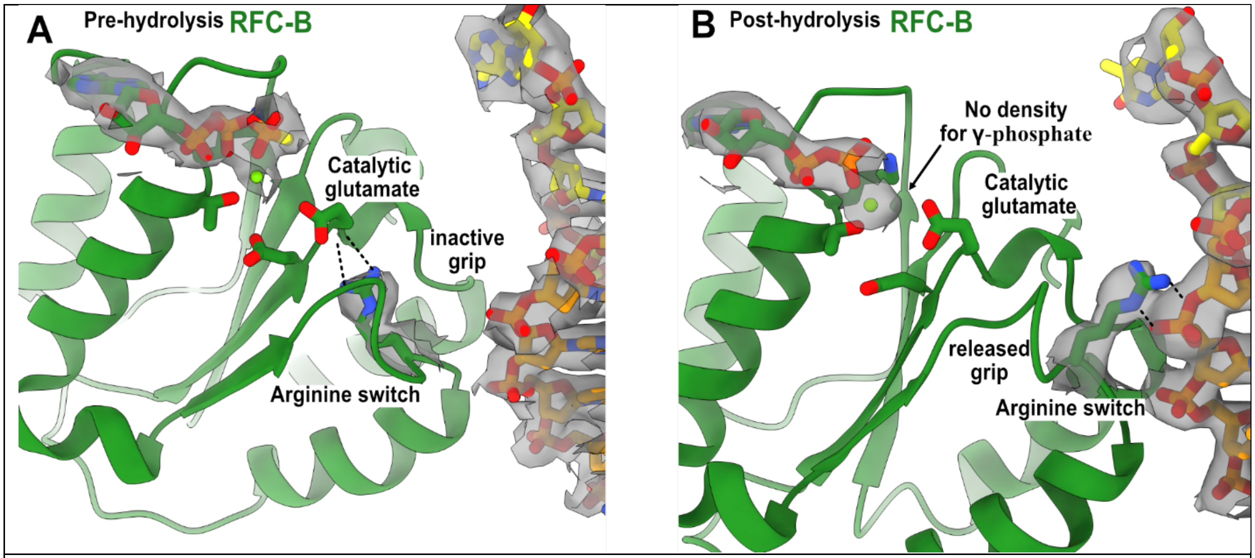


**Supplement Figure 6.2. Arginine switch residue is flipped into the active conformation.** **(A)** The active site of RFC-B is in pre-hydrolysis state in the structure of RFC:PCNA bound to p/t-DNA, and the conserved arginine switch residue holds the catalytic glutamate in an inactive conformation (PDB ID: 7TID) **(B)** Here, 5-gapped DNA is bound and the arginine switch instead interacts with DNA, releasing its grip from the catalytic glutamate. The catalytic glutamate is now considered to be in an active conformation. Accordingly, there is no density for the γ-phosphate of ATPγS in this conformation, indicating that hydrolysis has occurred at this site. We do not observe this hydrolysis in the structures bound to two p/t-DNAs or to nicked DNA, indicating that the presence of DNA at the external binding site is not driving ATP hydrolysis.
