## SupplementalTable1_map-model for "A second DNA binding site on RFC facilitates clamp loading at gapped or nicked DNA"

| Dataset | **RFC:PCNA with p/t DNA** | **RFC:PCNA with dsDNA with a 6 nucleotide gap** | **RFC:PCNA with dsDNA with a 5 nucleotide gap** | **RFC:PCNA with nicked DNA** |
|---|---|---|---|---|
| Magnification | 81,000 | 45,000 | 105,000 | 105,000 |
| Voltage (keV) | 300 | 200 | 300 | 300 |
| Cumulative exposure (e–/Å 2 ) | 40 | 48 | 48 | 49 |
| Detector | K3™ | K3™ | K3™ | K3™ |
| Pixel size (Å) | 1.06 | 0.87 | 0.83 | 0.83 |
| Defocus range (µm) | -1.2 to -2.3 | -1 to 2.2 | -1 to 2.2 | -1 to 2 |
| Micrographs used (no.) | 4499 | 4040 | 5118 | 4690 |
| Initial particle images (no.) | 1,331,440 | 797,499 | 1,098,517 | 874,202 |
| Symmetry | C1 | | | |
| **Class Name** | **RFC:PCNA bound to two p/t DNA molecules** | **RFC:PCNA bound to dsDNA with a 6 nucleotide gap** | **RFC:PCNA bound to dsDNA with a 5 nucleotide gap** | **RFC:PCNA bound to nicked DNA** |
| Final Refined particles (no.) | 43,129 | 130,421 | 271,745 | 119,631 |
| Map resolution (Å, FSC 0.143) | 3.4 | 3.3 | 3.0 | 3.7 |
| Model-Map CC_mask | nd | 0.82 | 0.81 | 0.8 |
| Bond lengths (Å), angles (°) | nd | 0.003, 0.628 | 0.002, 0.594 | 0.003, 0.687 |
| Ramachandran Outliers, Allowed, Favored | nd | 0.0, 2.13, 97.87 | 0.0, 1.64, 98.36 | 0.0, 1.98, 98.02 |
| Poor rotamers (%), MolProbity score, Clashscore (all atoms) | nd | 0.08, 1.66, 13.08 | 0.04, 1.56, 10.94 | 0.08, 1.64, 13.52 |
