## SupplementalTable2_DNA_sequences for "A second DNA binding site on RFC facilitates clamp loading at gapped or nicked DNA"

| Template | Sequence | primer | Sequence | Non primer | Sequence | Name used in assay |
|---|---|---|---|---|---|---|
| Template 30, T30 | TTTTTTTTTTTATGTACTCGTAGTGTCTGC | Primer20-2AP-0 | GCAGACACTACGAGTACAT/32AmPu/ | | | p/t-DNA p=0 |
| Template 30-T-1 | TTTTTTTTTTTTTGTACTCGTAGTGTCTGC-3’ | Primer20-2AP-1 | GCAGACACTACGAGTACA/i2AmPr/A | | | p/t-DNA p=-1 |
| Template 50-ap_gapped2 | TTGTGGGTAGATAAATACAGACCTAAGTCCTTTGTACTCGTAGTGTCTGC | Primer20-2AP-1 | GCAGACACTACGAGTACA/i2AmPr/A | 3' PrimerB24_gapped | AGGTCTGTATTTATCTACCCACAA | 6nt gap p=-1 |
| Same as above | | Same as above | | 3' PrimerB25_gapped | TAGGTCTGTATTTATCTACCCACAA | 5nt gap p=-1 |
| Same as above | | Same as above | | 3' PrimerB26_gapped | TTAGGTCTGTATTTATCTACCCACAA | 4nt gap p=-1 |
| Same as above | | Same as above | | 3'PrimerB30_gapped | GGACTTAGGTCTGTATTTATCTACCCACAA | Nicked p=-1 |
| Same as above | | Same as above | | 3'PrimerB30_gapped_P | /5Phos/GGACTTAGGTCTGTATTTATCTACCCACAA | Nicked 5’ PO4 p=-1 |
| Template 30-3'-T | TATGTACTCGTAGTGTCTGTTTTTTTTTTT | | | Primer20-2AP-20 | /52AmPr/CAGACACTACGAGTACATA | recessed 5' p=0 |

| | | | | | | |
|---|---|---|---|---|---|---|
| Template 50_gapped | TTGTGGGTAGATAAATACAGACCTAAGTCCTTGAATGCCGCGTGCGTCCC | 5'Primer 20_gapped | GGGACGCACGCGGCATTCAA | | | p/t-DNA |
| Same as above | | Same as above | | 3'PrimerB20_gapped | CTGTATTTATCTACCCACAA | 10 gap |
| Same as above | | 5'Primer 21_gapped | GGGACGCACGCGGCATTCAAG | Same as above | | 9 gap |
| Same as above | | 5'Primer 22_gapped | GGGACGCACGCGGCATTCAAGG | Same as above | | 8 gap |
| Same as above | | 5'Primer 23_gapped | GGGACGCACGCGGCATTCAAGGA | Same as above | | 7 gap |
| Same as above | | 5'Primer 24_gapped | GGGACGCACGCGGCATTCAAGGAC | Same as above | | 6 gap (used in ATPase assay and cryoEM) |
| Same as above | | 5'Primer 25_gapped | GGGACGCACGCGGCATTCAAGGACT | Same as above | | 5 gap (used in ATPase assay and cryoEM) |

| | | | | | | |
|---|---|---|---|---|---|---|
| Same as above | | 5'Primer 26_gapped | GGGACGCACGCGGCATTCAAGGACTT | Same as above | | 4 gap |
| Same as above | | 5'Primer 27_gapped | GGGACGCACGCGGCATTCAAGGACTTA | Same as above | | 3 gap |
| Same as above | | 5'Primer 28_gapped | GGGACGCACGCGGCATTCAAGGACTTAG | Same as above | | 2 gap |
| Same as above | | 5'Primer 29_gapped | GGGACGCACGCGGCATTCAAGGACTTAGG | Same as above | | 1 gap |
| Template 50-ap_gapped2 | TTGTGGGTAGATAAATACAGACCTAAGTCCTTTGTACTCGTAGTGTCTGC | Primer20-1 | GCAGACACTACGAGTACAAA | 3'PrimerB30_gapped_P | /5Phos/GGACTTAGGTCTGTATTTATCTACCCACAA | Nicked 5' PO4 (used in CryoEM) |
